# A thermostable phospholipase C obtained by consensus design

**DOI:** 10.1101/2022.10.06.511049

**Authors:** Diego S. Val, Luisina Di Nardo, Fiorela Marchisio, Salvador Peiru, María Eugenia Castelli, Luciano Abriata, Hugo G. Menzella, Rodolfo M. Rasia

## Abstract

Proteins’ extraordinary performance in recognition and catalysis have led their use in a range of applications. But proteins obtained from natural sources are oftentimes not suitable for direct use in industrial or diagnostic setups. Natural proteins, evolved to optimally perform a task in physiological conditions, usually lack the stability required to be used in harsher conditions. Therefore, the alteration of the stability of proteins is commonly pursued in protein engineering studies.

Here we achieved a substantial thermal stabilization of a bacterial Zn(II) dependent phospholipase C by consensus sequence design. We retrieved and analyzed sequenced homologs from different sources, selecting a subset of examples for expression and characterization. A non-natural consensus sequence showed the highest stability and activity among those tested. Comparison of activity and stability parameters of this stabilized mutant and other natural variants bearing similar mutations allow us to pinpoint the sites most likely to be responsible for the enhancement. Point mutations in these sites alter the unfolding process of the consensus sequence. We show that the stabilized version of the protein retains full activity even in the harsh oil degumming conditions, making it suitable for industrial applications.

---

Folding of a protein and the ensuing stability of its conformation is dictated by its amino acid sequence. There is a wealth of literature inquiring on the relationship between sequence, conformation and stability. While one can argue that the sequence-conformation relationship has been recently solved for globular proteins with the development of AlphaFold2 and similar models ^1^, the second part of the problem, that is predicting stability from the sequence (or even from an experimentally determined structure) remains largely unsolved. Therefore, engineering a protein to improve its thermodynamic stability is still an exceedingly difficult problem ^2^.

Over the last decades, several strategies for the stabilization of protein folds were proposed, ranging from structure-based rational mutagenesis to random mutagenesis and selection and, more recently, machine learning approaches ^3–5^. Among these, the use of a consensus sequence was suggested initially by Steipe *et al*. ^6^ and subsequently tested on several different proteins ^4,7–9^. This strategy takes advantage of the information contained in multiple sequence alignments of homologous proteins. Within a group of sequences corresponding to the same fold and function found in different organisms, neutral or destabilizing mutations are highly probable but have no selective pressure. In contrast, stabilizing mutations are less probable but will tend to be selected. Therefore, an artificial variant which sequence corresponds to the most probable amino acid at each position of a multiple sequence alignment is likely to result in a protein more stable than each of the individual proteins present in the alignment ^10^.

Variants of the phosphatidylcholine preferring *Bacillus cereus* phospholipase C are currently used in edible oil refining (Jiang et al.,2011). We have previously worked in the engineering of this enzyme (Elena et al., 2017; Ravasi et al., 2015), and in the search for natural thermostable variants (Marchisio et al., 2022). In this work we present the use of the consensus sequence approach to produce a new thermostable variant of this enzyme. We then investigate the basis of the enhanced thermostability, characterizing the kinetic features and folding thermodynamics of the consensus variant and four diverse natural PLCs. Finally we probe particular sites and their influence in the stability of the consensus variant. Application of this novel consensus PLC in vegetable oil refining will result in the broad adaptation of enzymatic degumming with a high economic and environmental impact^11^.

## Materials and methods

### Consensus sequence design and variant selection

A set of homologous sequences for *B. cereus* Zn(II) PLC was obtained using HHblits Alignment as provided by the Gremlin Server ^12^ (http://openseq.org/submit.php). The provided sequences were manually purged by keeping only those sequences showing all Zn(II) ligands present in *B. cereus* PLC: W1; H14; D55; H69; H118; D122; H128; H142 and E146 (*B. cereus* sequence numbering). The consensus sequence was then produced by selecting the most frequent residue present at each position (Supplementary Figure 1 ^13^).

We selected four natural sequences present in the starting alignment in order to express them and compare their performance with the consensus variant: *Bacillus cereus* PLC (CePLC), variants of which are used in industrial oil degumming; *Bacillus clarus* PLC (ClPLC), the natural sequence closest to the consensus; *Bacillus manliponensis* PLC (MaPLC), a sequence harboring a set of potentially stabilizing mutations present in the consensus but not in the closest homolog; *Bacillus pseudomycoides* PLC (PsPLC), a natural sequence closer to the *B. cereus* one (Supplementary Figure 1).

### Strains and growth conditions

The bacterial strains used in this study were *Escherichia coli* Top10, for general cloning purposes, and *Corynebacterium glutamicum* ATCC 13869, for protein expression. Both of them were cultivated in Luria-Bertani (LB) medium (tryptone, 10 g/L; yeast extract, 5 g/L; and NaCl 5 g/L) supplemented with 50 mg/L kanamycin with shaking at 200 rpm. Growth temperatures were 37 °C for *E. coli* and 30 °C for *C. glutamicum*.

### Sequence cloning and transformation

The CePLC expression plasmid used corresponds to that previously constructed in the group ^14^. The remaining sequences were codon-optimized and synthesized as described previously ^15^ and acquired from Twist Biosciences (CA, United States). The optimized sequences were designed with *Sac*II restriction site in the 5’ end and *Eco*RI in the 3’ end. Twist plasmids were purified using Axygen Biosciences AxyPrep™ Plasmid Miniprep Kit and digested with *Sac*II and *Eco*RI from New England Biolabs. The treated sequences were purified from 1.5 % agarose gel using Zymo Gene Clean up Recovery kit. The plasmid pTGR100 containing CePLC was treated with *Sac*II and *Eco*RI and the linear plasmid was purified. Ligation was performed with T4 DNA ligase from Thermofisher.

Preparation of competent cells and transformation of *E. coli* TOP10 and *C. glutamicum* ATCC 13869 were performed according to the protocols described ^16,17^.

### Protein production and purification

For the production of PLCs variants, C. glutamicum seeds carrying the expression vectors were grown overnight in LB medium. Then, a 100-fold dilution of the cultures was made and incubated with shaking at 30 °C until OD600 reached 0.5. At that moment, the cultures were induced with isopropyl-β-d-thiogalactopyranoside (IPTG, Genbiotech, Argentina) at a final concentration of 0.5 mM and left overnight with shaking.

The enzymes were recovered from the culture supernatants by precipitation with (NH4)2SO4, redissolved and incubated with Trypsin 0.5 ug/mL for 1h at 37 °C in order to obtain the mature proteins by cleavage of the propeptides, as previously described 14. Then, the proteins were loaded in a Shodex PH-814a hydrophobic interaction chromatography column and eluted with a (NH4)2SO4 gradient in Hepes 25 mM pH 7 on a HPLC system (Agilent 1200, USA). The recovered fractions were separated by SDS-PAGE on 12% gels, stained with Coomassie Brilliant blue to assess their purity and quantified by absorbance measurement at 280 nm 18.

### Thermal shift assay

To compare the thermal stability of the variants we employed the ThermoFluor method ^19^.

Basically, 5 ul of 1xSYPRO orange was added to a sample of each protein (45 ul of 5 uM) in 5 mM Hepes pH 7. The samples were heated from 25 to 95 °C at a rate of 1 °C per minute in PCR tubes. The fluorescence intensity was measured as a continuous function of temperature using a Veriti real-time PCR device (Applied Biosystems) to monitor protein unfolding. The data were normalized and analyzed using custom scripts in Python language.

### Activity measurements

PLC activity in aqueous buffer

The substrate 4-Nitrophenyl phosphorylcholine (NPPC, Toronto Research Chemicals) was used to measure the activity of enzymes in aqueous solution.

For time course reactions, 75 ul of NPPC 200 mM were added to a cuvette with 75 μl of 10 μM PLC. The reaction was monitored using a Jasco V-530 UV/VIS spectrophotometer until 80 % of the NPPC was consumed. These curves were used to determine the kinetic parameters of the different enzymes.

For stability experiments, the activity of the enzymes that were incubated at different temperatures was determined using an Epoch microplate reader (Biotek). In this case, 50 μl of a 10 μM enzyme sample were mixed with 50 μl of 2 mM NPPC. The reaction was followed for 10 minutes at 405 nm and the initial rate at each temperature was calculated.

### PLC activity in crude oil

Oil degumming experiments were conducted basically as previously described 20 using 5 mg of PLC per kg of oil. Concisely, 3 g of crude soybean oil (1200 ppm phosphate) was preincubated at the given temperature in a VP710 stirrer (V&P Scientic Inc.). Subsequently, enzymes were added and the mixture was incubated for 2 hours with continuous stirring.

For residual phosphate determination, degummed oil was treated with an Ultra-Turrax T8 homogenizer and 200 mg were mixed with 1 ml of 100 mM Tris-HCl pH 8.0. The mixture was incubated for 1 h at 37 °C with continuous stirring. Then, the mixture was centrifuged for 5 min at 14000 rpm. 200 ul of the aqueous phase were recovered and mixed with 200 ul of chloroform. After another centrifugation, 50 ul of aqueous solution were recovered and treated with 0.3 U of calf intestinal phosphatase (NEB) for 1 h at 37 °C. Finally, the solution was mixed with 440 μL of 5 % TCA and with 500 μL of color reagent (4 % FeSO4, 1 % (NH4)6MoO24·H2O, 3.2 % H2SO4), and absorbance recorded at 700 nm in a Epoch microplate reader. The concentration of phosphate was calculated from a standard curve containing 2.5 to 25 mM of inorganic phosphate.

### CD spectroscopy

Circular dichroism spectra were acquired on a Jasco J-1500 spectropolarimeter equipped with a Peltier temperature controlling unit. For far UV spectra we employed a 0.1 cm path cuvette to minimize the buffer contribution to absorption. Spectra were acquired from 210 to 240 nm, averaging four scans to improve the signal-to-noise ratio. The readout of the spectropolarimeter was converted to deg/cm^2^.dmol using the known concentration of protein and the average molecular weight of the amino acids. For near UV spectra we used a 1 cm micro cuvette acquiring data from 320 to 250 nm, averaging eight scans. Near UV CD spectra are informed in deg/mol.

### Evaluation of the stability of the variants

Enzyme samples were incubated in buffer at the selected temperature for different time steps. Subsequently the activity of the sample was measured using the chromogenic substrate as detailed in section 0. The decrease in activity with time was then fit to an exponential function to obtain the denaturation rate at the selected temperature. Finally, an Arrhenius plot (ln(k_obs_) vs. 1/T) was produced for each variant to obtain estimates for the activation energies.

## Results and discussion

With the aim of applying the consensus sequence approach to produce a stable variant of bacterial Zn(II) dependent phospholipase C we retrieved a set of natural homologs of CePLC. These 1059 natural sequences were filtered keeping only those harboring the known Zn(II) ligands to obtain the consensus sequence (here named ChPLC). This sequence and four other natural variants were codon optimized for expression in *C. glutamicum* ATCC 13869 and cloned in expression vectors to characterize their stability.

### Protein expression, spectroscopy and activity assays

We expressed all protein variants using *C. glutamicum* ATCC 13869 as expression host. All proteins were expressed at high levels and could be purified to homogeneity (Supplementary Figure 2).

To assess the fold conservation of the expressed proteins, we acquired far UV and near UV CD spectra (Figure 1). The far UV CD spectra of the variants are similar, showing that they all acquire the canonical helical fold of Zn(II) dependent bacterial PLCs. Near UV CD spectra are more dissimilar between the proteins, as expected due to the differences in aromatic amino acids content. Nevertheless, a significant signal intensity was obtained for all enzymes indicating the presence of structured aromatic residues in the fold and a stable tertiary structure.

**Figure 1.**
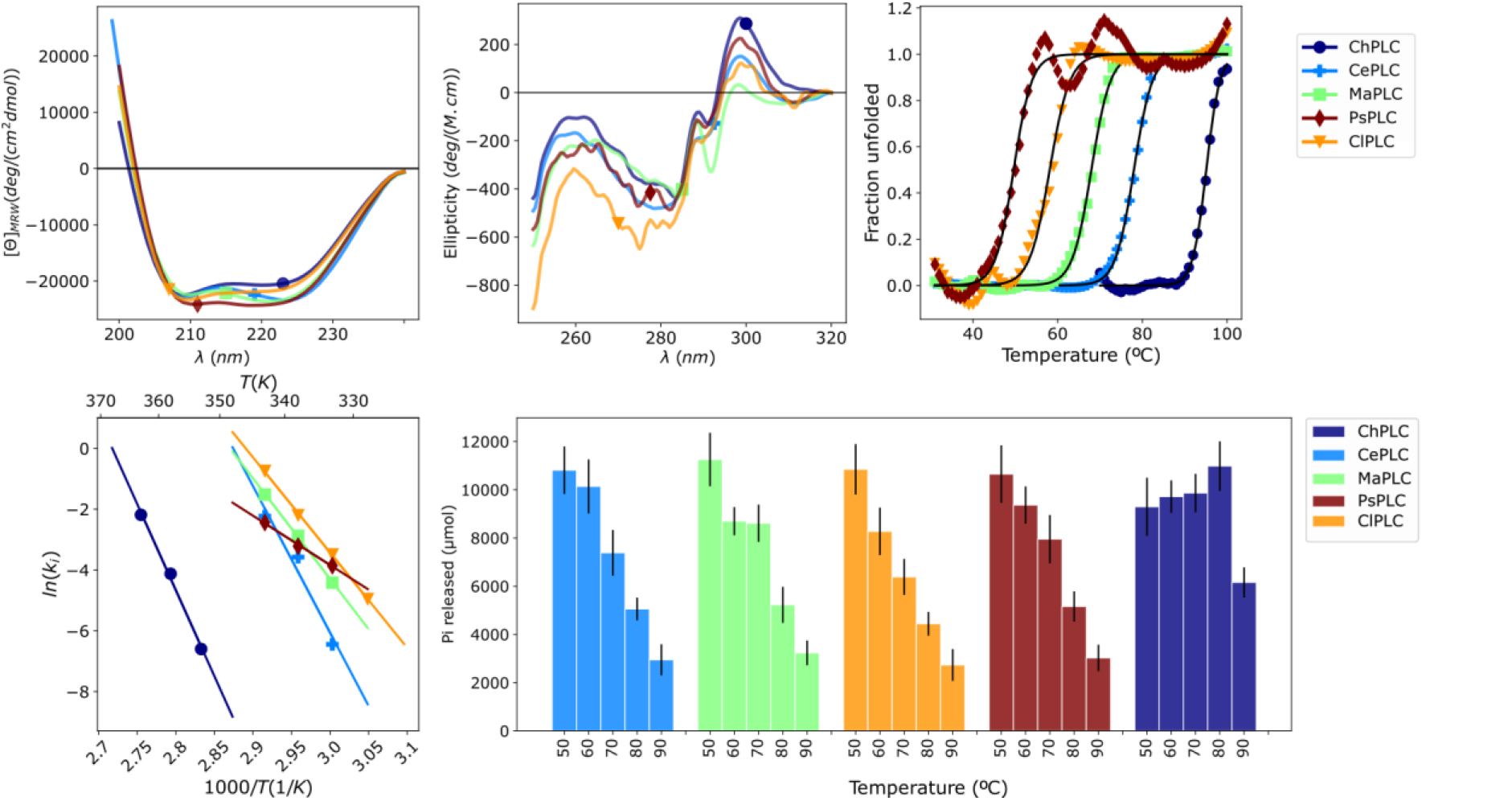
Biophysical and kinetic characterization of the enzyme variants tested. Top left, far UV CD spectra; top center, near UV CD spectra. Top right, thermal shift assay with Sypro Orange. Bottom left, Arrhenius plots showing the inactivation rate of the enzymes at different temperatures. Bottom right, activities in crude oil, expressed as amount of phosphate released, at different temperatures.

We then checked if the purified variants had indeed phospholipase C activity. We used the chromogenic substrate NPPC to measure enzymatic activity. Initial rate measurements of the reaction with CePLC performed up to 7 mM substrate, resulted in an almost linear dependence of the reaction rate with substrate concentration (Supplementary Figure 3). This indicates that the K_m_ of BcPLC for this substrate is much larger than 10 mM. We therefore decided to use low substrate concentrations so the reaction can be considered pseudo-first order in substrate with a rate constant equal to k_cat_/K_m_^21^. The obtained rate constants at 30 to 50 °C were similar for all variants, with the consensus protein showing the highest activity and the ClPLC and PsPLC variants showing the lowest activity (Table 1 and Supplementary Figure 3).

**Table 1.** k_cat_/K_m_, catalytic efficiency obtained at 40 °C using NPPC as substrate. Tm, denaturation temperatures measured by thermal shift assay. E^a^, activation energy for thermal denaturing measured by activity loss. Tm apoenzyme, melting temperature of the apoenzymes measured by far UV CD. Pi release, relative degumming activity at 80 vs 50 °C.

| Enzyme | k<sub>cat</sub>/K<sub>m</sub><br>(1/(M.s)) | T<sub>m</sub><br>(TSA, °C) | E<sup>a</sup><br>(kcal/mol) | T<sub>m</sub> apo<br>(°C) | K<sub>off</sub> Zn(II)<br>(1/min) | Pi release<br>80/50 °C |
| --- | --- | --- | --- | --- | --- | --- |
| ChPLC | 34.9 | 95.1 | 112.6 | 58.8 | 3.6 | 1.18 |
| CePLC | 30.8 | 78.0 | 96.0 | 40.7 | 3.1 | 0.47 |
| MaPLC | 14.5 | 68.0 | 65.9 | 58.4 | 3.1 | 0.46 |
| PsPLC | 15.9 | 49.6 | 57.6 | 49.9 | 4.0 | 0.48 |
| CIPLC | 23.3 | 58.1 | 62.4 | 38.4 | 8.3 | 0.41 |

### Evaluation of the stability of the different variants

We obtained an initial estimate of the stability of the different variants by means of a differential scanning fluorimetry assay using SYPRO Orange as dye (Figure 1). The transition temperatures range from 50 to 95 °C, increasing in order PsPLC < ClPLC < MaPLC < CePLC < ChPLC (Table 1). The melting point for the consensus variant as determined by this method is almost 20 °C higher than that of the more stable natural variant tested, CePLC.

Reaction time courses acquired between 55 and 80 °C show the same trend. In these experiments both substrate hydrolysis and enzyme inactivation occur simultaneously. ChPLC reaction rate increases continuously in the whole range, but the natural variants are inactivated as the temperature is raised (Supplementary Figure 4).

We then decided to quantify the relative stabilities by other methods. Both chemical (urea/GdmHCl) and thermal unfolding of mature CePLC are irreversible. For this reason, we resorted to the measurement of thermal unfolding kinetics to provide estimates for the activation energy (E^a^) of the irreversible denaturation step.

When protein unfolding is irreversible, incubation of enzyme samples at high temperature will result in a decay of the activity with a characteristic first order rate constant. The amount of active enzyme remaining in solution can then be estimated by measuring the residual activity at a lower temperature. Calculated first order rate constants at different temperatures can be subsequently used to obtain an estimate for the activation energy of the irreversible denaturation step ^22^. These energies are used in turn to compare quantitatively the relative stability of the different variants.

We obtained Arrhenius plots for all five variants tested (Figure 1). The estimated ΔG^++^_denat_ decrease in order ChPLC >> CePLC > MaPLC ≈ ClPLC > PsPLC (Table 1). The consensus variant displays a strikingly higher barrier for thermal denaturation. Unfolding of this enzyme requires 17 kcal/mole more energy than that of the more stable natural variant tested. More importantly, the stabilization brought about by the combination of all consensus residues, does not affect the catalytic efficiency of the enzyme, which is the highest among the variants tested in the present work.

The apo forms of bacterial Zn(II) PLCs are less stable than the Zn(II) bound forms, showing that Zn(II) binding in the active site contributes significantly to the overall stability of the folded protein ^23^. Therefore, the increased stability of the consensus enzyme could be due to an increase in Zn(II) binding affinity. We determined the rate of Zn(II) loss of the variants by measuring the rate of the activity loss in the presence of the metal chelator EDTA (Supplementary Figure 4), but found no significant differences between them, ruling out this hypothesis.

Finally, we evaluated the relative stabilities of the apoenzymes with a temperature scan followed by far UV CD (Supplementary Figure 4). We found that both ChPLC and MaPLC show similar stabilities in the apo-form, with melting temperatures higher than the other variants.

### Activity in crude oil

Phospholipases C are employed in enzymatic degumming of edible oils. The reaction conditions in this process are radically different from the reaction in aqueous solution. The enzyme functions in a lamellar liquid-crystalline phase formed by water, oil and the phospholipids that act as surfactant ^24,25^. This environment could change significantly the stability parameters observed in aqueous solution.

We therefore tested if the stability of the enzymes in the conditions of the degumming process followed the same trend as the stability measured in buffer. We measured the amount of inorganic phosphate released after a two-hour degumming reaction at temperatures ranging from 50 to 90 °C with 10 °C intervals (Figure 1 and Table 1). All natural variants show a decrease in the amount of phosphate released with increasing temperature, down to ca. 1/3 of the maximum amount of phosphate released at 90 °C. In contrast, the consensus variant shows a slight increase in phosphate released up to 80 °C, lowering the efficiency at 90 °C but still releasing twice as much phosphate as the any of the other natural variants at this temperature. The consensus variant showing the highest activity in the degumming assay at 80 °C makes it extraordinarily valuable for edible oil refinement ^11^.

### Cooperative effect of consensus residues

We modeled the structures of all variants produced using AlphaFold2 in order to find a rationale for the stability of the consensus protein (Figure 2). Supplementary Table 1 shows the interatomic contacts calculated using the Arpeggio web server ^26^. This analysis shows that both ChPLC and MaPLC have a higher number of aromatic and hydrophobic contacts than the other variants. These two variants display the highest melting temperature in the apo form. We therefore analyzed in more detail the environment of aromatic residues in the modeled structure of ChPLC.

**Figure 2.**
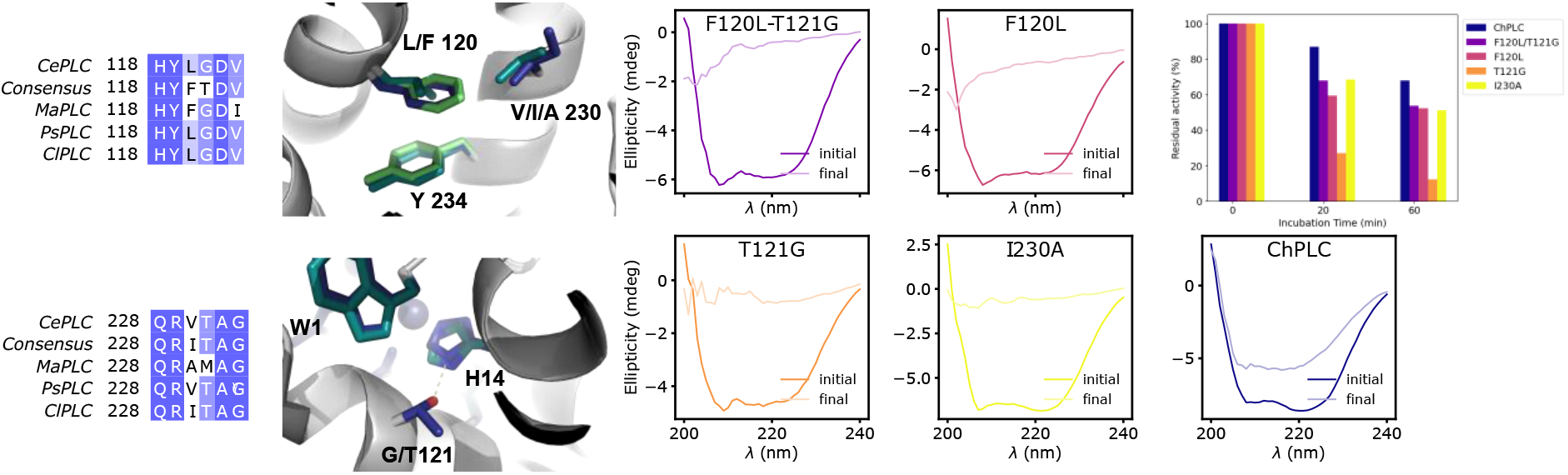
Left, highlights of structural features at the base of the stability increase of ChPLC. Regions of the structure of CePLC (PDB 1AH7) and the AlphaFold2 modeled structures of ChPLC and MaPLC are overlaid with the sidechain carbon atoms color coded as follows: CePLC, cyan; MaPLC, light green; ChPLC, dark blue. Right, far UV CD spectra of the apo ChPLC mutants before and after a temperature ramp from 20 to 95 ºC. Top right, residual activity after incubation at 65 ºC.

The phenylalanine at position 120 can induce a significant stabilization to the consensus protein adding an aromatic stacking interaction with the conserved tyrosine 234. Among the variants tested, only the MaPLC sequence shows the F120-Y234 aromatic stacking. In this natural variant, the large sidechain volume of F120 appears compensated by the reduction in volume of the opposite residue 230, with an alanine substituting a larger hydrophobic residue prevalent in the other sequences (Figure 2). In contrast, the consensus sequence shows an isoleucine at position 230, but the large sidechain appears to pack in the core without distorting the structure.

The larger increase in E^a^ and melting temperature found for holo ChPLC indicates that the Zn(II) site contributes significantly to the stability of this variant. When analyzing the second shell ligand residues in the active site, we found that ChPLC introduces a threonine in position 121 where all other variants tested have a glycine. The T121 sidechain can H-bond with the sidechain of H14, one of the Zn(II) ligands, potentially increasing the stability of the Zn(II) site and thus of the protein as a whole.

Based on these observations, we generated three point mutants within the consensus sequence: F120L, T121G, and I230A. Additionally, we produced the double mutant F120A/T121G. As anticipated, all mutant variants exhibited activity against the chromogenic substrate and a proper folding, as indicated by their CD spectra (Figure 2). The mutations had negligible impact on the stability determined through thermal shift assays. The melting temperatures of the apo forms closely resembled that of ChPLC (Supplementary Figure 5). Interestingly, while apo ChPLC experienced partially reversible temperature-induced unfolding, the mutants did not recover their secondary structure following the assay (Figure 2). The melting transition of apo ChPLC followed by far UV CD was also shallower, suggesting a slower unfolding process. All mutants show a discrete transition in the HT values of the temperature scan, coincident with the CD transition, probably reflecting an aggregation induced increase of the optical density. In contrast, the HT values for ChPLC increase smoothly along the temperature ramp (Supplementary Figure 5), suggesting that aggregation is not taking place. To delve deeper, we assessed the unfolding kinetics of the apo forms at 65°C. This process occurred in two distinct phases, with the faster phase taking place within the experimental dead time. Notably, all mutants displayed faster unfolding rates, with a larger amplitude in the fast phase compared to ChPLC (Supplementary Figure 5). Lastly, we evaluated the stability of all mutants by measuring residual activity after incubation at increasing temperatures. All mutations led to a decrease in stability over time when compared with ChPLC (Figure 2).

In summary, our analysis of ChPLC’s point mutants reveals a collective synergy among the evaluated sites, contributing significantly to its exceptional stability. This stability arises not only from an elevated energy barrier hindering unfolding but also from a degree of reversibility within the unfolding process. Notably, the engineered consensus variant surpasses all natural sequences and point mutants tested in this study in terms of stability, while retaining full enzymatic activity. We anticipate that this enhanced enzyme will find valuable applications within the edible oil industry.^11^

## Supporting information

supp_figures_ACS_Biochemistry

## Supporting Information

Sequences used to obtain the consensus; PAGE of purified PLC variants; Kinetic characterization of the PLC variants; Zn(II) release and stability of the apo forms; Stability of the mutant forms; Interatomic interactions in the variants (PDF).

## Acknowledgements

This work was supported by grants PICT-2016-4124; PICT-2019-01528 and IO 1027-00332 to RMR.

DV and LDN are recipients of fellowships from CONICET. Add other acknowledgments here

## Notes

### Competing Interest Statement

The authors have declared no competing interest.

### Summary of Updates

The manuscript has been revised to include a thermal characterization of a set of point mutants in the thermostable enzyme, ChPLC, which is described in this paper. This characterization aims to scan and explain the enzyme's thermosensitivity.

