## Supplementary material for "A thermostable phospholipase C obtained by consensus design": supp_figures_ACS_Biochemistry

**Supplementary Figure 1.** Alignment of the sequences used to obtain the consensus.

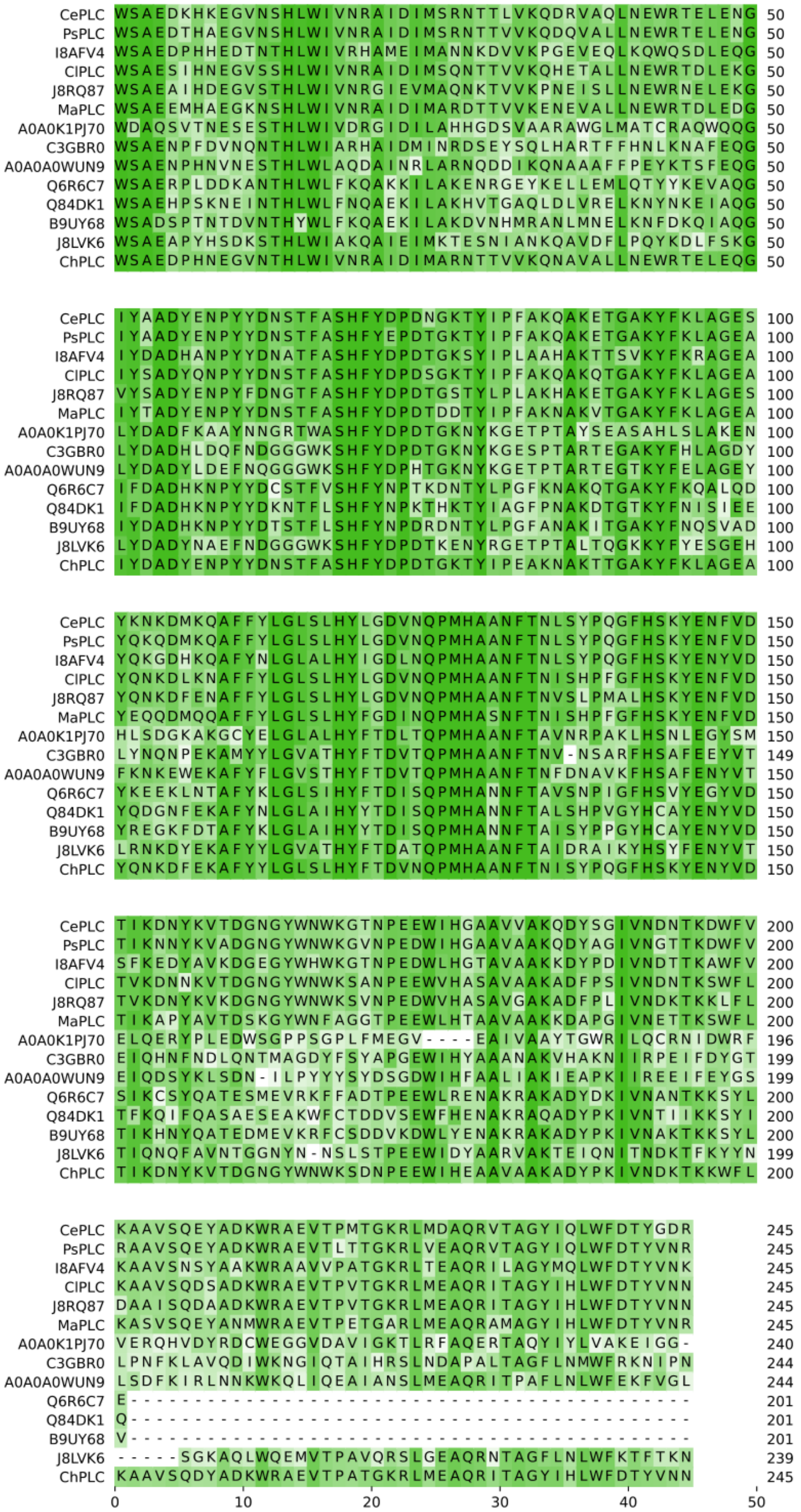

**Supplementary Figure 2.** Expression and purification of the PLC variants. For each protein, the pro-enzyme and the activated enzyme, with the propeptide separated by trypsin digestion, are loaded side by side. Arrows indicate the mature proteins. PPS: Prestained Protein Standard, broad range (10-200 kDa).

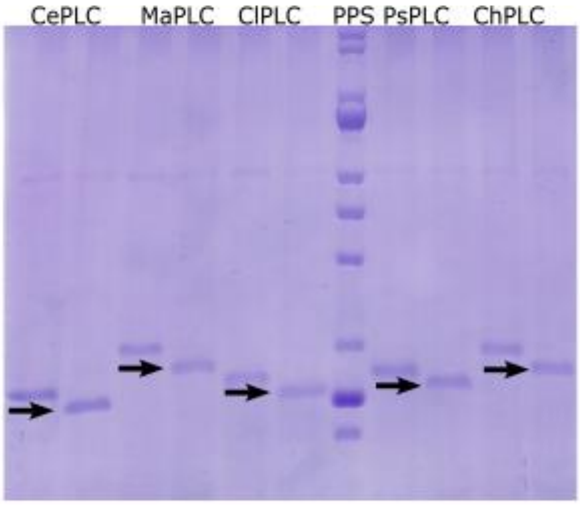

**Supplementary Figure 3.** Kinetics of PLC reactions in aqueous buffer with NPPC. A, initial rates of the reaction of CePLC with NPPC. B-D, reaction time courses of 100 µM NPPC with each of the enzymes at (B) 30 ºC; (C) 40 ºC, (D) 50 ºC. Fit of the data points to an exponential function are shown as black lines. E, kcat/KM values obtained from the exponential fits. (F) Inactivation of the enzymes during reaction time courses with NPPC at different temperatures.

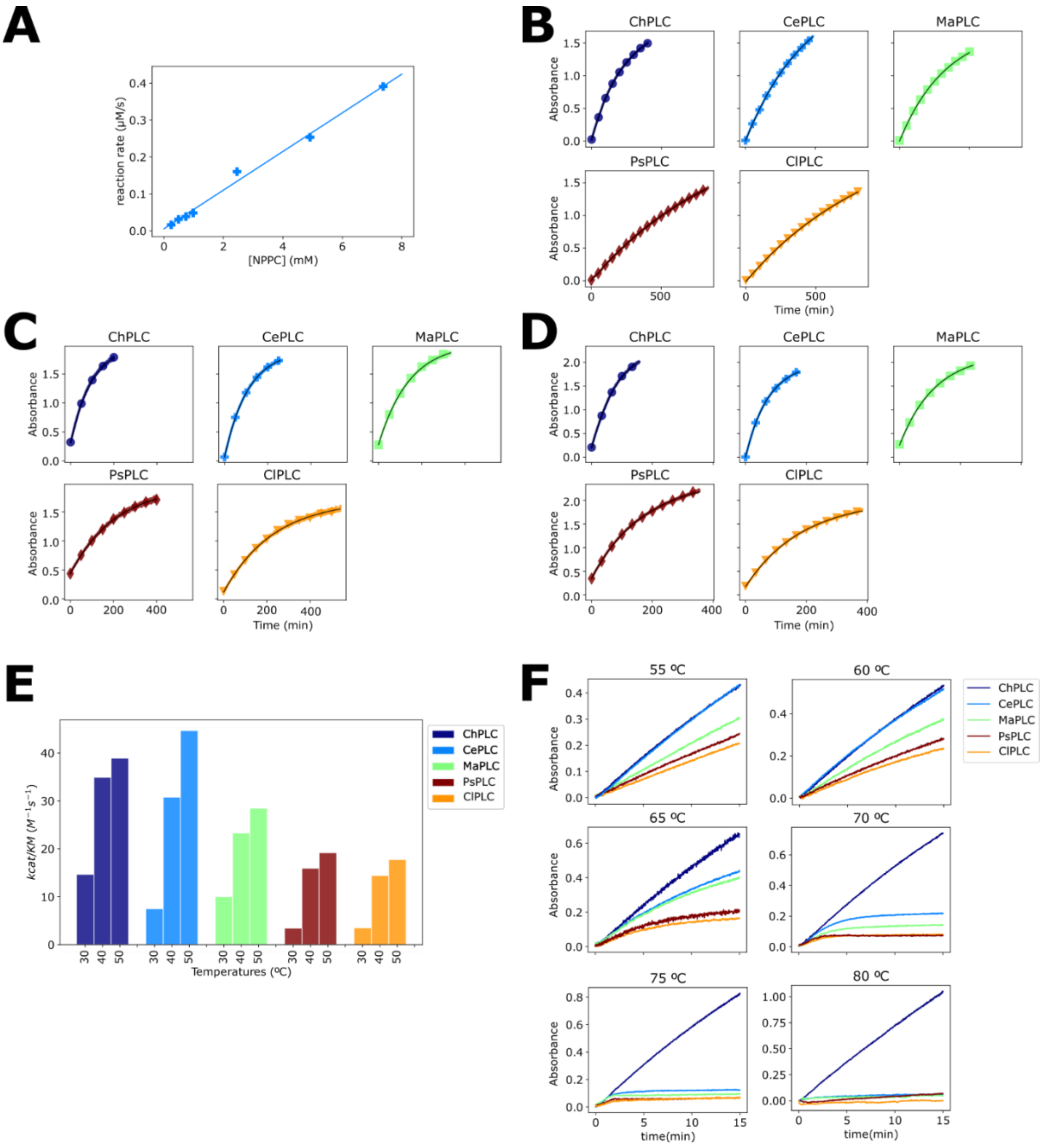

**Supplementary Figure 4. A.** Left, kinetics of Zn(II) loss, as determined by the enzymatic activity remaining in time after incubation with 5 mM EDTA. Each column is a repetition of the experiment. B, structural stability of the apoenzymes followed by CD at 222 nm with temperature scanning. Pre- and post-transition signal region were fit to first order polynomials and the whole curves normalized to obtain the fraction of unfolded protein. The curves are offset to improve visibility.

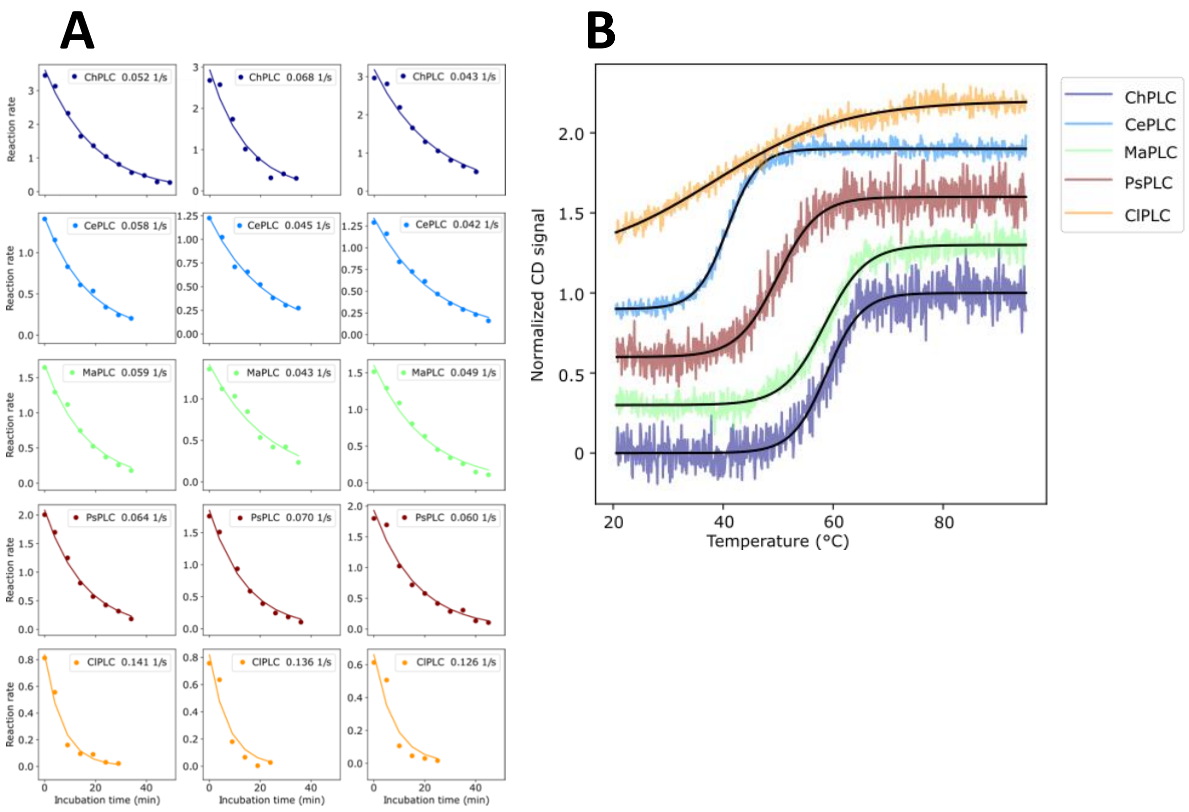

**Supplementary Figure 5.** Stability of ChPLC mutants. A, unfolding of the apo forms followed by far UV-CD. B. Photomultiplier voltages recorded during the experiments shown in A. C. Unfolding kinetics of the mutant proteins at 65 ºC followed by CD at 222 nm.

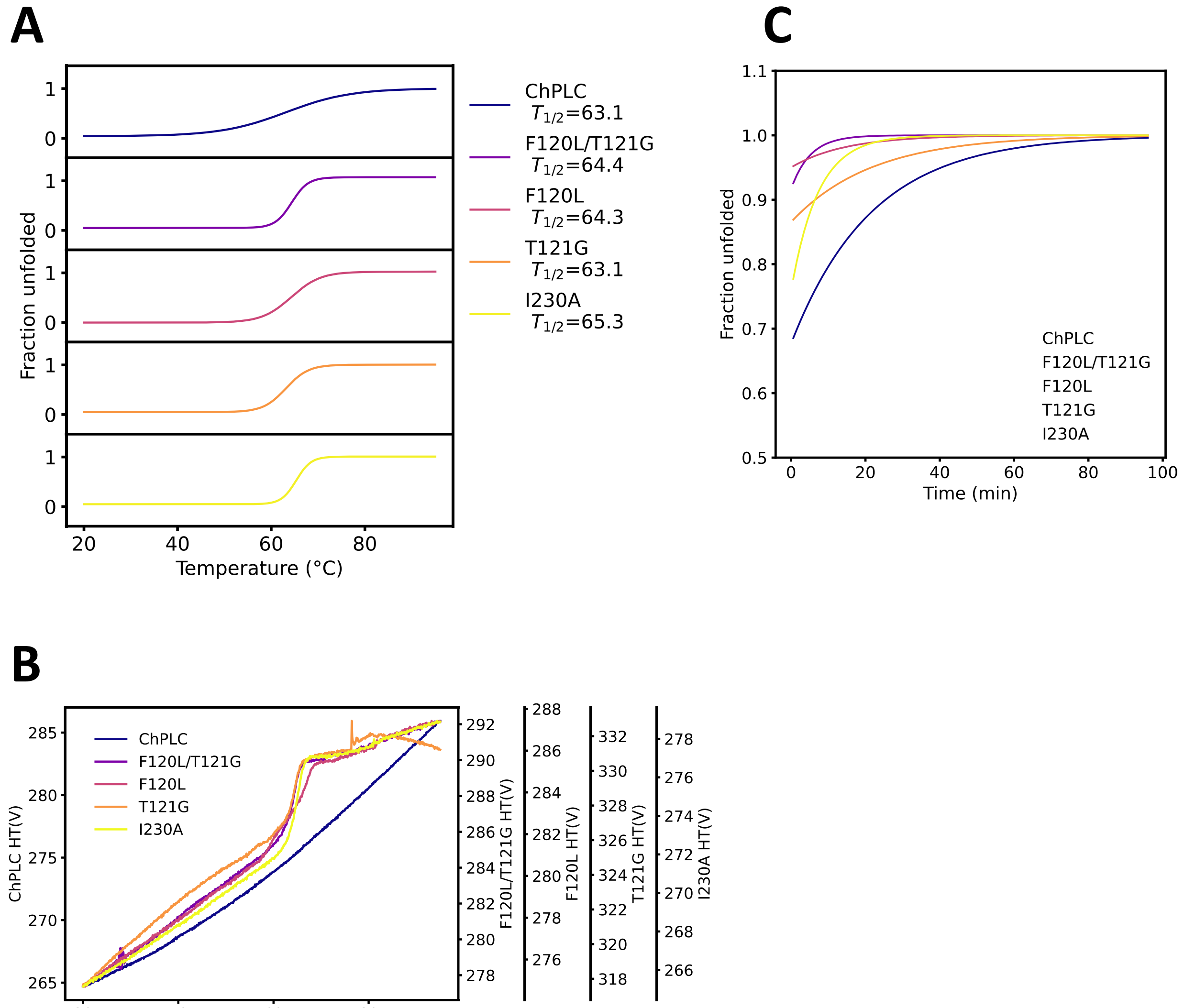

**Supplementary Table 1.** Interatomic interactions calculated for the five variants characterized. The enzymes are sorted by melting temperature of the apo forms.

| | Weak Hbond | Hbond | Ionic | Aromatic | Hydrophobic | Carbonyl | Polar | Weak Polar | $T_m$ apo (ºC) |
|---|---|---|---|---|---|---|---|---|---|
| **ChPLC** | 329 | 222 | 71 | 105 | 739 | 8 | 525 | 286 | 58.8 |
| **MaPLC** | 342 | 218 | 65 | 98 | 739 | 12 | 572 | 262 | 58.4 |
| **PsPLC** | 342 | 209 | 47 | 89 | 691 | 11 | 577 | 250 | 49.9 |
| **CePLC** | 338 | 226 | 55 | 88 | 692 | 10 | 575 | 254 | 40.7 |
| **ClPLC** | 336 | 224 | 66 | 82 | 679 | 10 | 585 | 259 | 38.4 |
